# Rotavirus re-programs multiple IFN receptors and restricts their intestinal antiviral and inflammatory functions

**DOI:** 10.1101/702837

**Authors:** Adrish Sen, Nima D. Namsa, Ningguo Feng, Harry B. Greenberg

**Affiliations:** The Department of Microbiology and Immunology, Stanford University School of Medicine, Stanford, California, USA; VA Palo Alto Health Care System, Palo Alto, California, USA; Department of Molecular Biology and Biotechnology, Tezpur University (A Central University), Napaam 784 028, Assam, India

## Abstract

Rotaviruses (RV) cause acute severe diarrhea in the absence of substantial intestinal inflammation. They are also highly infectious in their homologous host species. The efficient replication capacity of RV in the small bowel is substantially linked to its ability to inhibit different types of interferons (IFNs). Here, we find that during RV infection *in vitro*, both virus-infected and uninfected bystander cells resist STAT1 phosphorylation and IRF7 induction in response to exogenous IFN. Functionally, cellular transcription in response to exogenous stimulation with IFN, but not intracellular dsRNA, was inhibited by RV. Further, IFNAR1 stimulation during RV infection significantly repressed a set of virus-induced transcripts. Regulation of IFN signaling *in vivo* was studied in suckling mice using the highly infectious homologous murine EW RV strain. Kinetic studies indicated that whereas sustained EW RV replication and IFN induction occurred in the small intestine, IFN-stimulated transcripts significantly decreased over time. In addition, LPS-mediated intestinal damage, driven by STAT1-induced inflammation, was prevented in EW RV-infected mice. Remarkably, ectopic stimulation of either IFNAR1 or IFNGR1 in murine RV-infected mice eliminated several intestinal antiviral and inflammatory transcriptional responses to RV. In contrast to homologous murine RV, infection with a STAT1-sensitive heterologous simian RV strain induced multiple IFN-stimulated transcripts, inflammatory cytokines, and intestinal expression of STAT1-pY701. Finally, RV strain-specific STAT1 regulation in the gut plays a prominent role in the activation of multiple intestinal caspases. On the other hand, the simian RRV strain, but not murine EW RV, uniquely triggers the cleavage of both extrinsic and intrinsic caspases (−8, −9, and −3) in a STAT1-mediated manner. Collectively, these findings reveal efficient re-programing of multiple IFN receptors in the gut towards a negative feedback mode of signaling, accompanied by suppression of IFN-mediated antiviral, apoptotic, and inflammatory functions, during natural RV intestinal infection.

## INTRODUCTION

Rotaviruses are non-enveloped segmented dsRNA viruses that cause severe dehydrating diarrhea in many mammals and an estimated 200,000 human infant deaths annually. Acute RV infection in a homologous host (i.e. the host species in which a specific RV strain replicates to high titer and spreads from individual to individual) is an extremely efficient process; severe diarrhea can be initiated by exposure to a very small infectious inoculum^1^. The host innate interferon (IFN) response is a multi-pronged line of defense against many viral pathogens as its induction can activate hundreds of different antiviral genes and trigger inflammatory signals – thus eliminating infected cells, limiting the spread of virus infection to uninfected bystander cells, and shaping both the magnitude and quality of subsequent adaptive immune responses^2^.

All three major IFN types (α/β, γ, and λ) are induced during homologous RV infection of the small bowel^3,4^, but their antiviral effects on viral replication are modest at best. Although relatively poorly understood, the inflammatory effects of IFNs are also likely impaired during RV infection, which is associated with only mild intestinal inflammation compared to other acute diarrheal pathogens^5–7^. Previously, in a suckling mouse model of RV disease, we demonstrated that the IFNAR1, IFNGR1, and IFNLR1 receptors, as well as the downstream convergent transcription factor STAT1, play an important role in restricting heterologous RV (RV not typically isolated from the symptomatically infected host species) replication^1,3,4,8,9^. Depletion of IFNRs in suckling mice also results in an acute biliary inflammatory disease (often lethal) following infection with a heterologous (rhesus RRV) but not a homologous (murine EW RV) strain^8,10,11^. How homologous RV efficiently restricts the ability of different IFNs to execute their antiviral and inflammatory programs is not well understood but important both for more effective strategies to combat RV-associated mortalities and to better understand the varying pathogenic potentials of naturally circulating RV strains.

## MATERIALS AND METHODS

### Viruses and reagents

The tissue culture-adapted simian RRV and porcine SB1-A RV strains were propagated in African green monkey kidney cells (MA104) as previously described^12^. Virus titers were determined by plaque assays in MA104 cells. HT-29 cells, a colon cancer derived cell line with a variety of small bowel features^13^, were plated in 6-well or 24-well cluster plates and completely confluent monolayers were infected 2-3d later as described^14^. The non-tissue culture-adapted murine EW RV strain was prepared as an intestinal homogenate from RV-infected Sv129 suckling mice. The diarrheal dose (DD_50_) was determined as described^9^. Human intestinal epithelial HT29 and embryonic kidney HEK293 cells were purchased from the American Type Culture Collection (ATCC) and maintained in Advanced D-MEM Ham’s F12 (Cellgro) or D-MEM medium (D-MEM, Cellgro) containing 10% fetal calf serum (Invitrogen) supplemented with non-essential amino acids, glutamine, penicillin, and streptomycin, respectively. Purified human IFN-β (PBL) was used for stimulation of cells. Purified carrier-free universal type I IFN A/D (PBL), murine IFN-γ (PBL), LPS (Sigma) were used in animal studies. High-molecular weight LyoVec poly(I:C) (Invivogen) was used at 2.5 μg/mL.

### Mouse infection and tissue harvest

All mice were housed in the VA Palo Alto Health Care System (VAPAHCS) Veterinary Medicine Unit and all experimentation followed VAPAHCS Institutional Animal Care and Use Committee approved protocols. 3-5 day old Sv129 pups were infected via oral gavage with 10^4^ DD_50_ of murine EW RV, 4 × 10^6^ PFU of the simian RRV, or mock infected as described^9^. At the indicated time-points, the small intestines were removed. Sections were collected in Trizol for RNA purification or in 2× Laemmli buffer for protein analysis by immunoblotting, as described^4^ and stored at −80 °C, or fixed in 4% paraformaldehyde for processing and subsequent analyses as described^15^.

### Mouse stimulation and Luminex assays

To ectopically stimulate IFNAR1 and IFNGR1, suckling mice infected with 10^4^ DD_50_ of murine EW RV (or mock-infected) for 12 hours were intraperitoneally administered purified murine IFN-γ (1μg IFN-γ per pup) or universal type I IFN A/D (1μg per pup) and sacrificed 12 h later for analysis of transcripts by qRT-PCR as described^4^. For endotoxin stimulation, 3-5 day old 129sv mice were orally inoculated by gavage with 10^4^ DD_50_ of murine RV-EW. At 3 dpi, mice were intraperitoneally injected with PBS or purified LPS (Sigma, 10 mg/kg body weight) and sacrificed 6 h later for analysis of paraformaldehyde-fixed and paraffin embedded small bowel tissue sections by hematoxylin and eosin staining. For measurement of intestinal cytokines, mouse small intestines were harvested and weighed, and pooled tissues (from 3 pups) were homogenized in ice-cold PBS containing a cocktail of protease and phosphatase inhibitors (Sigma) using a handheld micropestle (Thermo Fisher). Lysates were clarified by two sequential rounds of centrifugation at 10,000× G for 10 min, and analyzed for cytokine levels at the Human Immune Monitoring Center at Stanford University. Mouse 38-plex kits were purchased from eBiosciences/Affymetrix and used according to the manufacturer’s recommendations. Plates were read using a Luminex 200 instrument with a lower bound of 50 beads per sample per cytokine. Custom assay Control beads by Radix Biosolutions are added to all wells. Murine IFN-β was measured using a commercial ELISA kit as per the manufacturer’s instructions (PBL).

### Western blot analysis

Preparation of cell lysates and immunoblotting *in vitro* was performed as described^12^. For mouse samples, intestinal tissue pieces were lysed in radioimmunoprecipitation assay buffer (RIPA) supplemented with protease and phosphatase inhibitors (Roche). Equal volumes of 2× Laemmli buffer (Sigma-Aldrich) were added to each sample. Viscosity was reduced by passing repeatedly through a 22G syringe and samples were boiled for 5 minutes prior to analysis by SDS-PAGE and immunoblotting. Blots were probed with primary antibodies directed towards the following antigens: STAT1-pY701, cleaved caspase 8, caspase 9, cleaved caspase 3, (Cell Signaling Technologies), RV VP6 (Santa Cruz Biotechnologies), and B-Actin (Sigma). The blots were subsequently incubated with either anti-rabbit or anti-mouse secondary antibodies conjugated to horseradish peroxidase (HRP, Amersham), incubated with ECL detection reagent, and exposed to film (GE Healthcare). All densitometry quantifications of protein blots were performed using ImageJ software analysis and normalized to actin protein bands.

### Transcriptional analysis

For *in vi*v*o* analysis of transcripts, mice were sacrificed and sections of the small intestine were collected on ice in Trizol (Life Technologies). For *in vitro* studies, cells were first washed in PBS and then lysed in Trizol. Total RNA was extracted following the manufacturer’s instructions and subjected to DNAse digestion before use in qRT-PCR. Synthesis of cDNA and subsequent microfluidics PCR on the Fluidigm platform was done as described earlier^16^ using commercially available taqman assays (Thermo-Fisher) and RV NSP5-specific assays described earlier^3,4,9^. Serial 10-fold dilutions of mouse or human reference RNA (Agilent) and blank controls were run in duplicate for each PCR run.

### Flow cytometry analysis

Staining and flow cytometry analysis of HT-29 cells was performed as described^14^. Cells were plated in 24-well cluster plates and infected with RV. Infected cells were harvested at 6 hpi or 16 hpi and transfected cells at 48 after transfection for flow cytometry analysis. Cells were washed in PBS and fixed at room temperature for 10 min using 1.6% (v/v) methanol-free paraformaldehyde (Electron Microscopy Services). Cells were washed in FACS staining buffer (PBS containing 0.5% w/v bovine serum albumin) and permeabilized in cold methanol at 4 °C for 10min. Cells were washed in FACS staining buffer, stained using antibodies directed to either STAT1-pY701 or IRF7 (from BD Biosciences), or to the RV VP6 antigen^14^ at 4 °C for 30 min, and washed prior to analysis by flow cytometry on a LSRII instrument (BD Biosciences). The flow data was analyzed using FlowJo software.

### Statistical analysis

Analysis of variance (ANOVA) was carried out by Kruskal-Wallis method using Dunn correction for multiple comparisons. Protein bands from SDS-PAGE blots were analyzed using Holm-Sidak t-test. P values <0.05 were considered significant as indicated in the figure legends. All error bars represent the standard error of the mean.

## RESULTS

### Rotavirus restricts IFN-directed STAT1 activation and IRF7 expression

Prior studies using several RV strains identified two distinct RV strategies to regulate IFN induction that are exemplified by either porcine (degrades β-TRCP) or simian (degrades IRFs) RV strains^17^. In contrast to this distinction, we reported that both porcine SB1-A and simian RRV strains similarly mediated the lysosomal degradation of endogenous IFNAR1, IFNGR1, and IFNLR1 subunits in RV infected (RV+) cells^15^ and prevented IFN-directed STAT1 phosphorylation at the Y701 epitope^14^, which is a critical convergent signal transduction event downstream of the different IFNRs. Surprisingly, in other *in vitro* studies, SB1-A was also found to inhibit IFN-directed STAT1-Y701 phosphorylation in RV- bystander cells, which expressed normal levels of IFN receptors^14^. Here, we extended our studies by examining whether IFN signaling is also inhibited in RV- bystander cells by the simian RV strain RRV. HT-29 cells were infected with RRV at a multiplicity of infection resulting in approximately 35% RV infected VP6+ cells (Fig. 1A, column 1) and at 12 hpi cells were stimulated for 2h with increasing doses of purified universal type I IFN (IFN-A/D) to induce STAT1-Y701 phosphorylation (Fig. 1A, columns 2-6). Analysis of RV VP6 and STAT1-pY701 expression by FACS demonstrated that mock-infected controls underwent a dose-dependent increase in STAT1-pY701 expression in response to ectopic IFN (Fig. 1A, row 1 and Fig. 1B). In contrast, both RRV VP6+ (RV+) and RRV VP6− (bystander) cell populations effectively resisted IFN-stimulated STAT1 phosphorylation, even at the highest concentration of IFN-A/D tested (Fig. 1A, row 2 and Fig. 1B). These findings demonstrate that the IFN signaling pathway is inhibited in uninfected bystander cells *in vitro* by the simian RV (RRV), similar to our earlier findings using a porcine RV^14^. Thus, although uninfected RV bystander cells express normal levels of IFNRs^15^ during infection, their ability to transduce IFN-directed signals is significantly impaired.

**Figure 1.**
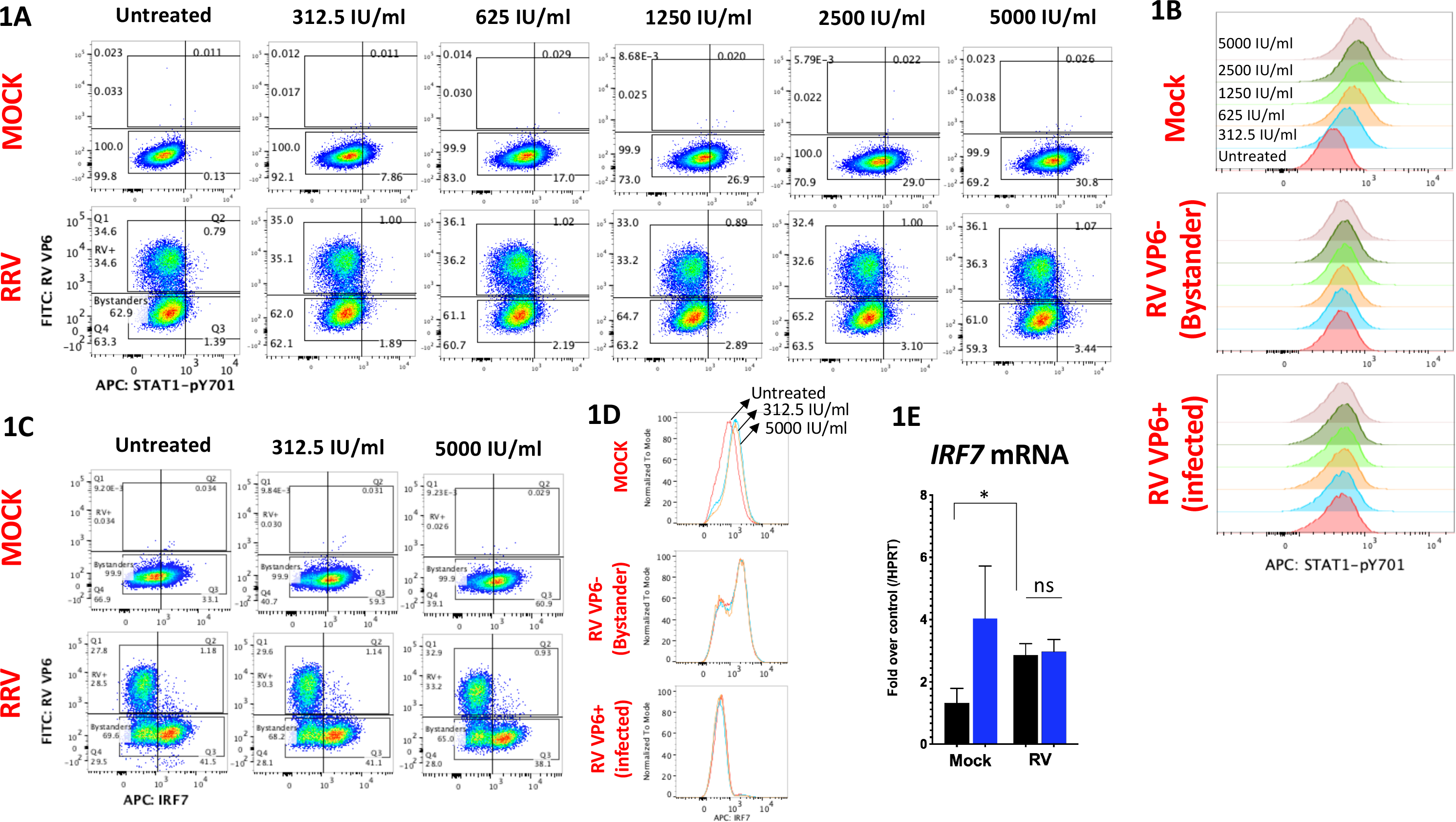
Rotavirus inhibits IFN-stimulated cellular responses *in vitro*. (A-B) HT-29 cells were infected with RRV (or mock infected) at a MOI of 1. At 10 hpi cells were stimulated with purified IFN-β at the indicated concentrations for 2 hours and analyzed by flow cytometry. (C-D) cells were infected with RRV and analyzed by FACS as in (A). (E) Cells infected in triplicate as in (C) and at 12 hpi were stimulated with PBS (black bars) or 500 IU/ml of purified IFN-β (blue bars) for 6h before analysis by qRT-PCR for IRF7 and HPRT transcript levels. (*p<0.02, *ns* not significant). Data shown are representative of two or more independent experiments.

To extend these observations, we examined the effect of simian RRV strain on levels of endogenous IRF7, an IFN-stimulated protein that is also targeted for degradation by the RRV-encoded NSP1 protein^18^. Analysis of RV VP6 and IRF7 by FACS was performed in HT-29 cells as described above (Fig. 1C). In mock controls, IRF7 was constitutively expressed by ~33% cells even in the absence of IFN stimulation. Following stimulation with either a low (312.5 IU/ml) or high (5,000 IU/ml) dose of IFN-A/D, a similar increase in IRF7 expression occurred (~60% cells were IRF7+ post-IFN stimulation at either dosage) (Fig. 1C, row 1 and Fig. 1D), indicating that relatively low IFN concentrations are sufficient to maximally induce IRF7 expression in HT-29 cells. Following RV infection, RRV VP6− cells exhibited elevated basal IRF7 expression in the absence of any exogenous IFN stimulation (i.e. ~43% of all cells, almost entirely deriving from the VP6− gate, were IRF7+), which could be possibly triggered by low levels of IFN induced earlier during RV infection^12,19^. However, IFN stimulation failed to induce any further increase in IRF7 expression in RRV VP6− cells (Fig. 1C, row 2 and Fig. 1D). In contrast, RV+ HT-29 cells expressed substantially lower levels of IRF7 (only ~4% VP6+ cells were IRF7+). Notably, no detectable increase in IRF7 expression occurred in infected cells following IFN-A/D stimulation at either dose. Overall, in these cell monolayers the percentage of HT-29 cells expressing IRF7 after stimulation with 5,000 IU/ml IFN was lower during RRV infection (~39% of all cells) than mock infection (~61% of all cells). By qRT-PCR analysis, we confirmed that although RRV infection induced IRF7 transcription compared to mock controls, it prevented any further exogenous IFN-stimulated IRF7 transcription (Fig. 1E). These initial data demonstrate that RV-infected cells are highly resistant to IFN-mediated induction of IRF7 or STAT1-pY701. In addition, the findings indicate that although RRV induces IRF7 in bystander cells during infection, it effectively prevents further IRF7 induction by saturating doses of IFN stimulus, which is in general agreement with the observed blockage of STAT1 phosphorylation in RV bystander cells (Fig. 1A). Since optimal IFN-directed STAT1-pY701 and IRF7 induction are important for IFNR positive feedforward transcriptional amplification^20^, it seemed likely that RV infection might perturb this process in both infected and bystander cells.

### Rotavirus perturbs interferon-stimulated feedforward cellular transcriptional response

To gain further insight into the regulation of the IFN signaling response by RV, we next examined the ability of ectopic IFN to upregulate antiviral and inflammatory transcription following RRV infection of HT-29 cells. Since several ISGs and inflammatory transcripts upregulated during IFN signaling are also induced by intracellular dsRNA^21^, primarily through NF-kB and IRF3 signaling pathways, we compared cellular transcriptional responses to either exogenously applied IFN-β or intracellular long dsRNA introduced by transfection. Following RV infection for 6h, cells were stimulated with either purified IFN-β or with liposomally-complexed long dsRNA for an additional 6h and then analyzed by qRT-PCR. The results (Fig. 2A) demonstrate that mock-infected HT-29 cells underwent significant upregulation of several transcripts in response to either IFN and/or dsRNA stimulation (*IFIT3*, *IFI6*, *CCL5*, *TNF*, *HERC5* were induced by both types of stimuli; *SERPING*, *CXCL10*, *IFI16*, *TNFAIP3* by IFN-A/D only; *IL29*, *CCL3*, *IFNG* by dsRNA only). In RV-infected samples, significant transcriptional upregulation of *CCL5*, *TNF*, *IL29*, *CCL3*, and *IRF8* occurred only in response to intracellular dsRNA. In addition, *RSAD2*, *IFIT3*, *IFI6*, *CXCL10*, *IFI16* and *TNFAIP3* transcripts were also weakly, but not significantly, elevated in response to dsRNA during RV infection. In contrast, we failed to detect any significant upregulation of measured transcripts in response to IFN-β with the sole exception of *SERPING*, whose induction was weaker in RV-infected samples than in mock-infected controls (Fig. 2A). The stimulus-specific RV effect on cellular transcription was unlikely due to differences in the level of RV replication, since it was unaltered by either IFN-β or dsRNA stimulation (Fig. 2B).

**Figure 2.**
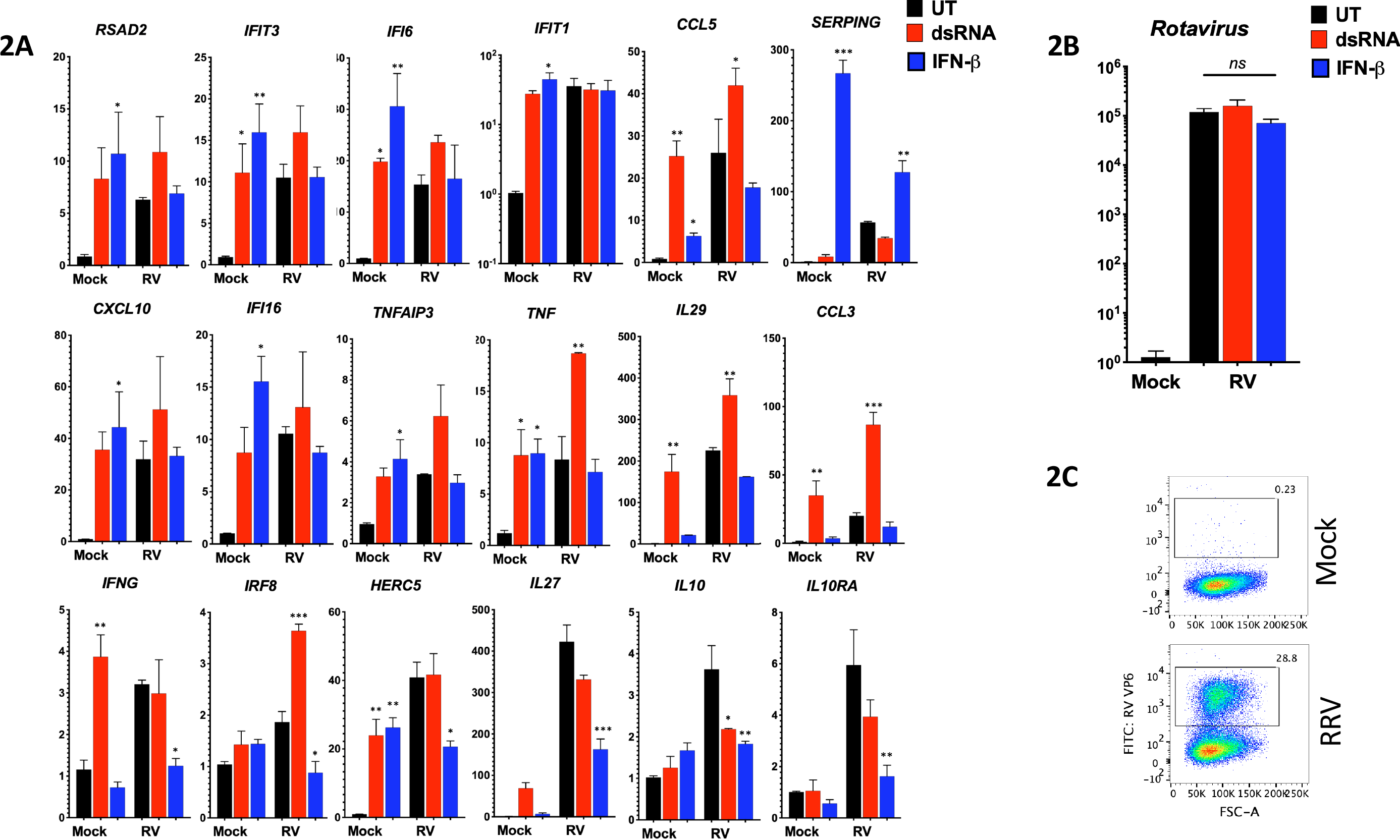
Rotavirus infection inhibits IFN-stimulated feedforward cellular transcription. (A-B) HT-29 cells grown in triplicate wells were infected with RRV strain at a MOI of 1 (or mock-infected) for 6h and then stimulated with 2μg of [>200 bp long dsRNA:liposome] complexes, or 500 IU/ml of purified IFN-β. At 12 hpi cells were lysed for RNA purification and then analyzed by qRT-PCR for expression of indicated host (A) and RV (B) transcripts. (*ns*, not significant, **\***p<0.02, **p<0.002, ***p<0.0002; significance is shown relative to untreated samples in mock and RV infected groups). (C) Cells were infected as in (A), and the percentage of cells expressing RV VP6 antigen before lysis was determined by FACS. Box indicates gates used to estimate RV-infected cells in the monolayer). Data shown is representative of two experiments.

Interestingly, ectopic stimulation of IFNAR1 during RRV infection also significantly repressed the expression of a set of transcripts from their levels in the unstimulated RRV samples (*IFNG*, *IRF8*, *HERC5*, *IL27*, *IL10*, and *IL10RA*) (Fig. 2A, bottom row). Although RV infection completely prevented (or repressed) the IFN-directed upregulation of transcripts, by FACS analysis we found that only ~30% of the cells in the monolayer were RRV VP6+ (Fig. 2C). Hence, these results suggested that RV inhibits IFN-directed cellular transcription in both RV-infected and in uninfected bystander cells.

### Rotavirus infection also restricts intestinal IFN amplification responses *in vivo*

The transcriptional amplification of IFNs (e.g. *IL28*, *IFNG*), antiviral genes (e,g, *MX2*), and inflammatory cytokines (e.g. *CXCL10*) occurs via STAT1 activation as part of the IFN feedforward signaling circuit^22^. We next examined the kinetics of selected IFN stimulated transcripts in the small intestine of suckling mice infected with a highly infectious homologous murine EW RV strain^1^ for either 12h or 72h (Fig. 3A-B). This analysis revealed that *IFNA4* (a primarily induced IFN gene, i.e. prior to the IFN amplification phase^*20*^) and *RV NSP3* transcripts continued to significantly increase between 12-72 hpi. In contrast, a significant decrease in RV-induced intestinal type II and III IFNs (*IL28* and *IFNG*), *RSAD2, IFIT2*, and *CXCL10* transcription occurred in EW RV-infected mice between 12-72 hpi (Fig. 3A). RV-induced transcription of *MX2*, a specific marker of the intestinal epithelial response to IFN stimulation^23^, also significantly declined from 12 to 72 hpi. These results indicate that during murine EW RV infection of suckling mice, despite sustained viral replication and IFN induction, several intestinal IFN amplification responses are down-regulated by 72 hpi. To further examine this finding, we directly tested whether intestinal damage caused by endotoxin, which mediates inflammation in a STAT1-dependent manner^24^, is also down-regulated by prior infection with EW RV. Suckling mice orally infected with EW RV (or mock) for 72h were systemically administered endotoxin and resulting intestinal damage assessed 6h later. In mock-infected pups, parenteral administration of endotoxin led to severe intestinal necrosis and tissue destruction (Fig. 3C). In contrast, suckling mice that were infected with EW RV prior to endotoxin challenge exhibited minimal small intestinal damage (Fig. 3C). These results further indicate that RV infection in the small bowel can suppress intestinal IFN-associated amplification responses.

**Figure 3.**
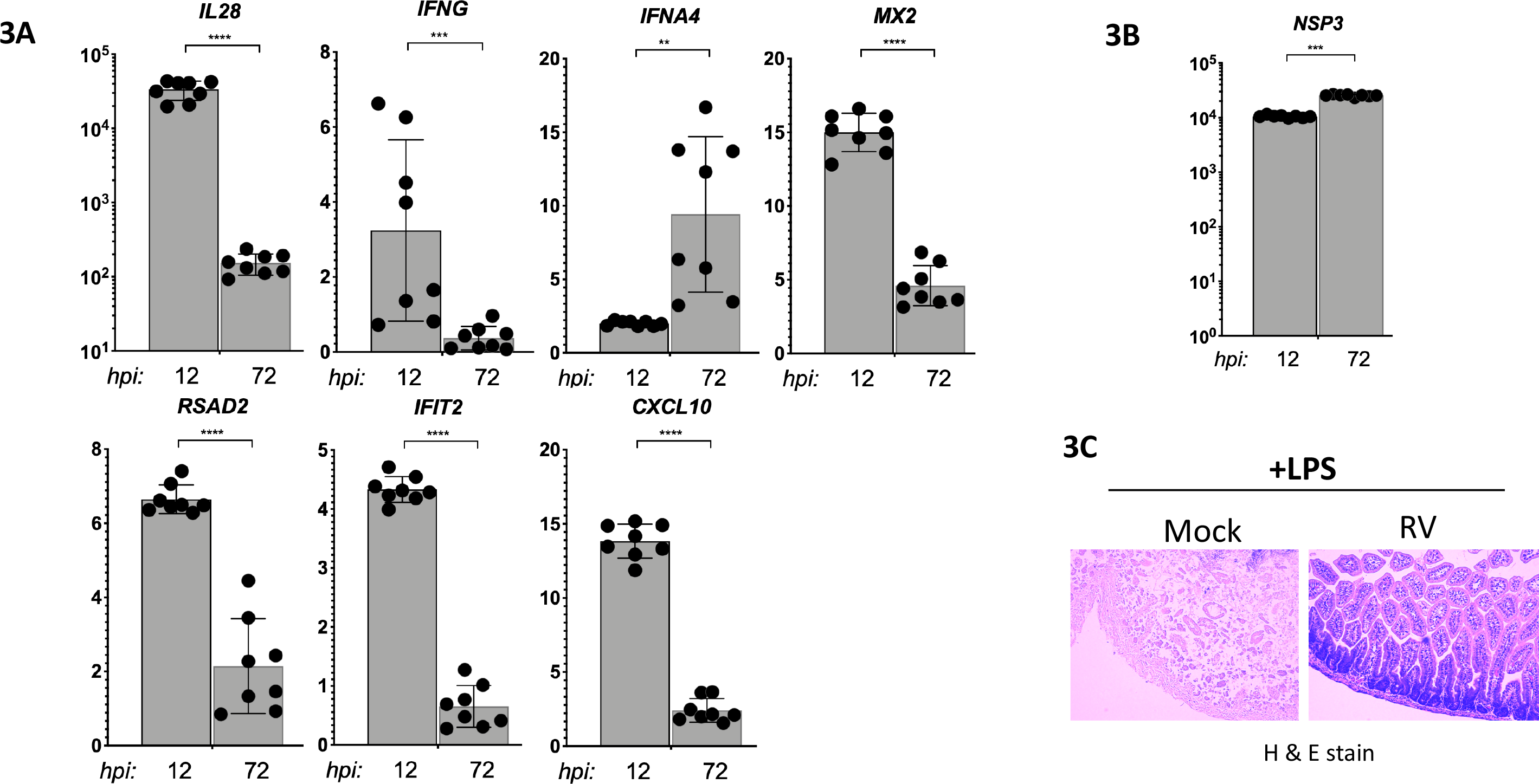
Effects of homologous RV infection on intestinal IFN amplification responses. (A-B) Suckling mice were infected with murine EW RV (or were mock infected, N=3-8 mice per group) for the times indicated, and small intestines collected to analyze the expression of transcripts shown. Data shown is fold change in EW RV versus mock infected mice. (C) Suckling mice were mock- or EW RV-infected and 3 d later were parenterally administered purified endotoxin (15μg/pup) for 6 h before harvesting small intestines for histological examination by hematoxylin and eosin staining. Data shown is representative of 2 pups per group.

### Rotavirus reprograms intestinal IFN type I and II receptors towards negative feedback signaling

Our data *in vitro* (Fig. 2A) indicated that ectopic IFN-stimulated transcription was suppressed during RV infection. Therefore, we next examined intestinal transcriptional responses to ectopically administered type I and type II IFNs in suckling mice infected with murine RV. Following EW RV infection for 12h, pups were parenterally administered either purified IFN-A/D or IFN-γ (or PBS), and small intestines were collected after an additional 12h for transcriptional qRT-PCR analysis (Fig. 4A). In the absence of any ectopic IFN stimulation, RV infection transcriptionally induced several intestinal IFNs, ISGs, and inflammatory genes, as expected from our prior findings (Fig. 3A). Remarkably, ectopic stimulation of IFNAR1 or IFNGR1 in the murine RV infected pups resulted in either a significant suppression (*IL28*, *IFIH1*) or the complete elimination (*IFNB1*, *IFNA5*, *IFIT1*, *MX2*, *TNFSF10*, *CXCL10*, *STAT1*) of several innate transcriptional responses to the RV infection. A similar IFN-dependent effect was not observed for *PDCD1*, *PCNA*, *GAPDH*, or *IKBKB* transcripts, or for *ARG2* and *IL12B*, whose expression during RV infection significantly increased following administration of either type of IFN (Fig. 4A). To ascertain whether the suppressive effect of IFN stimulation on RV associated transcription could be explained by the exogenous IFN mediated inhibition of EW RV replication, we measured intestinal RV RNA by qRT-PCR using two different EW RV-specific PCR assays (Fig. 4B). These results demonstrated that EW RV replication was not significantly altered following either IFN-A/D or IFN-γ stimulation under these conditions (Fig. 4B). These results reveal that during RV infection, exogenous stimulation of either type I and II IFNRs in the small intestine has the potential to efficiently suppress both antiviral and inflammatory transcriptional responses.

**Figure 4.**
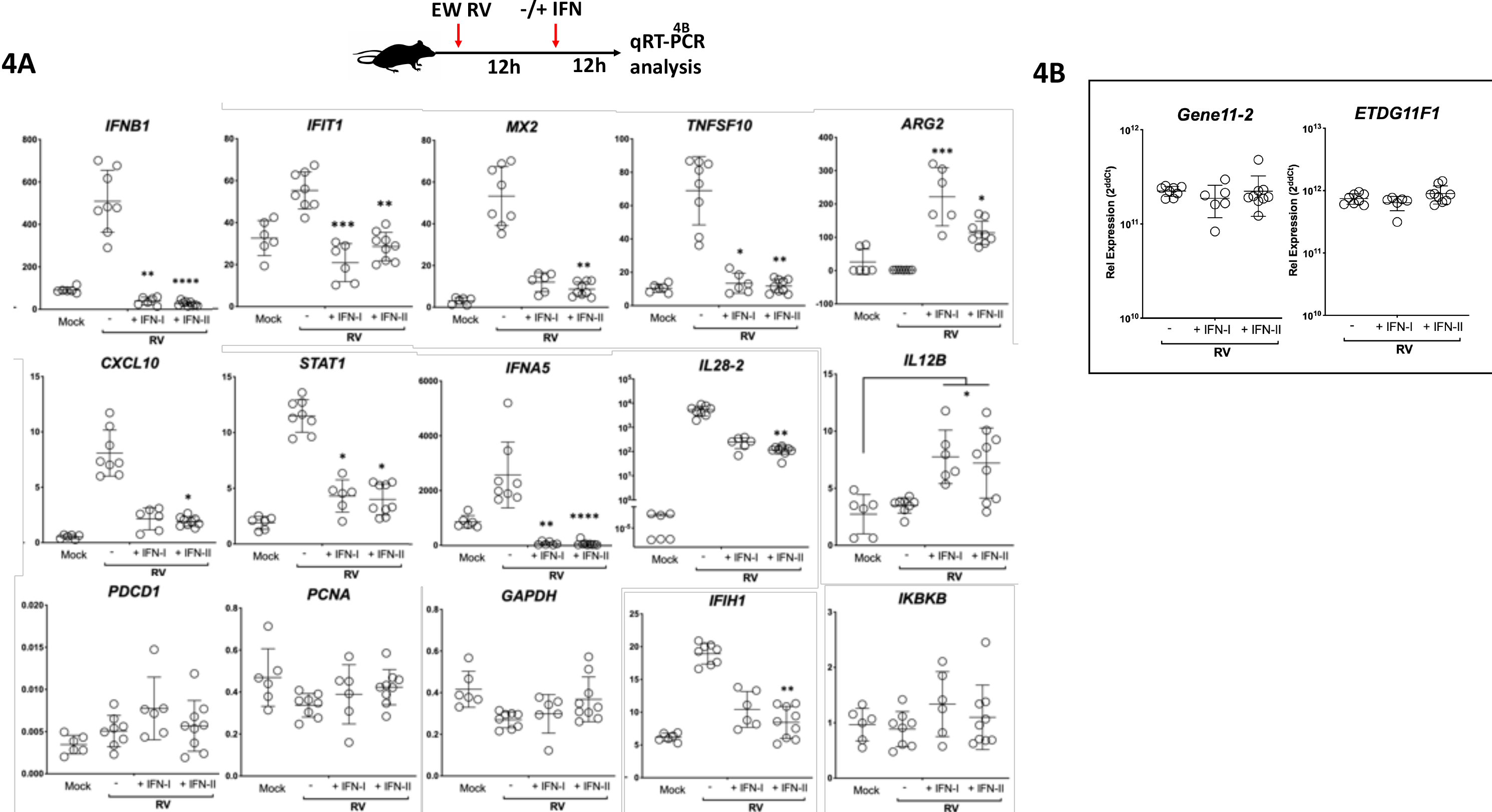
Rotavirus reprograms intestinal IFNAR1 and IFNGR1 towards a negative feedback mode of transcription. Schematic of the experimental approach used is shown at the top. Mice infected with EW RV (or mock) for 12 h were administered purified universal IFN-A/D (IFN-I), murine IFNγ (IFN-II), or PBS by the i.p. route. Twelve hours later mice were sacrificed, and small intestinal RNA analyzed by qRT-PCR for different host (A) or RV gene (B) transcripts. (**\***p<0.02, **p<0.002, ***p<0.0002; significance in each panel is relative to PBS-treated EW RV infected mice, unless indicated otherwise). Data is measured from 3-5 pups per group.

### Intestinal IFN signaling is regulated differently during infection with homologous versus heterologous RV*s*

The finding that murine EW RV impairs IFN signaling in suckling mice is consistent with studies in mice^8^ and human enteroids^25^ that demonstrate RV replication in the homologous host species is IFN-resistant. In contrast, we previously observed that replication of the heterologous simian RRV strain in suckling mice is highly sensitive to different types of IFN and the presence of STAT1 (by 2-4 logs)^3,4,8,9^. We thus compared intestinal IFN-stimulated antiviral genes and cytokines in mice infected with the heterologous simian RRV, homologous murine EW, or mock-infected. As shown in Figure 5A, at 12 hpi the intestinal replication of heterologous RRV was substantially and significantly lower than the homologous EW RV. However, RRV infection triggered significantly higher transcription levels of the ISG20, IFI203, and PELI1 genes as compared to EW RV (Fig. 5A). In addition, heterologous RV infection also resulted in significantly elevated intestinal protein levels of IFN-β and the IFN-stimulated pro-inflammatory cytokines IP10 and RANTES when compared to homologous RV (Fig. 5B). The expression of type III IFN (IL28) was also elevated in RRV-infected pups compared to either the EW RV or mock groups.

**Figure 5.**
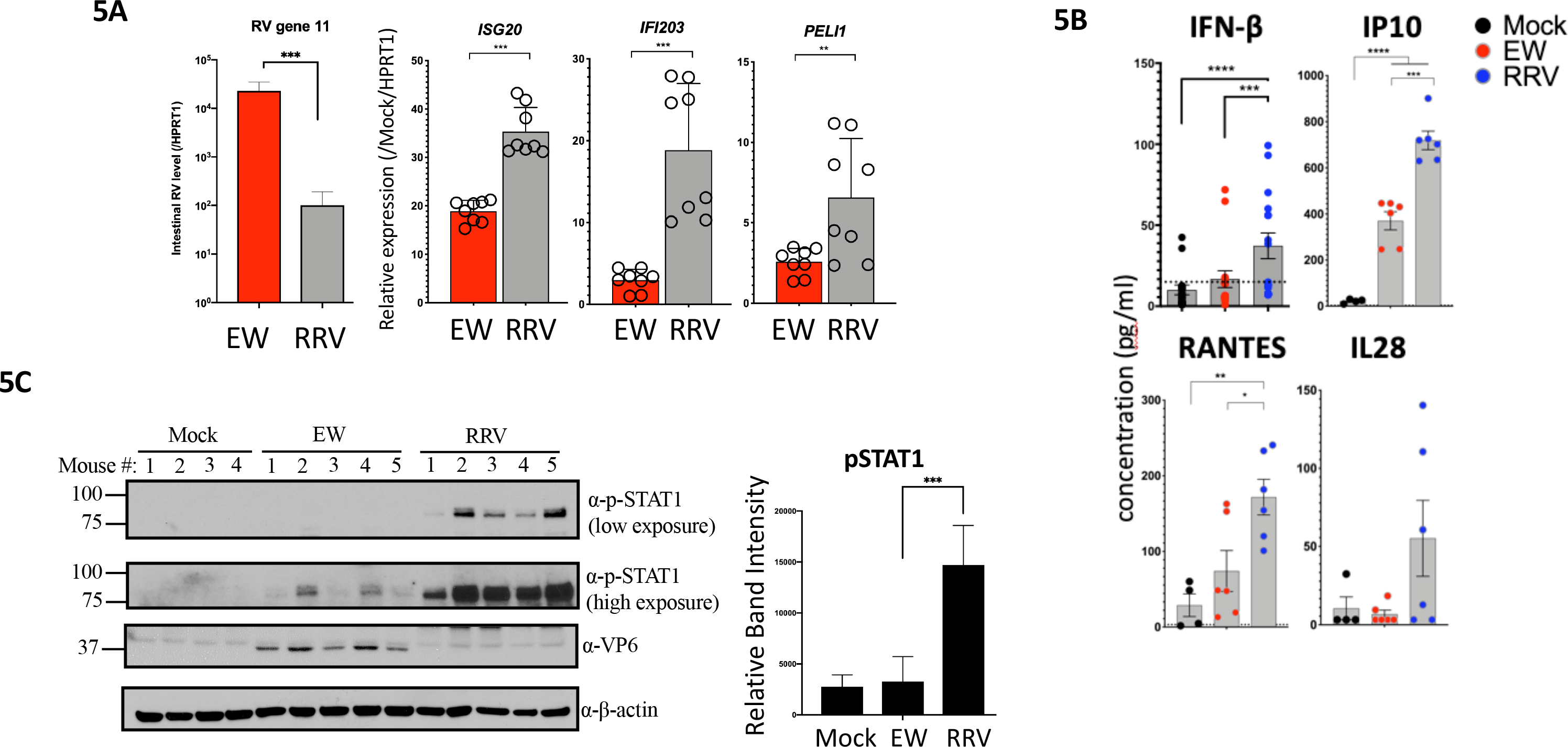
Differences between homologous and heterologous RV strains on IFN stimulated genes, inflammatory cytokine expression, and STAT1-Y701 activation *in vivo*. (A) Suckling mice (N= 3-5 pups per group) were infected with homologous murine EW or heterologous simian RRV RV strains (or mock infected). At 12 hpi mice were sacrificed for subsequent analysis of small intestinal transcripts by qRT-PCR. Data shown is fold relative to mock-infected mice. (B) Mice infected as in (A) were sacrificed and small intestinal biopsies were analyzed for the expression of the proteins shown either by ELISA (IFN-β) or Luminex-based cytokine assays. (C) Mice infected as in (A) were sacrificed and small intestines analyzed for immunoblotting analysis. Densitometric quantitation of pSTAT1 bands is shown in the bar diagram at the right. (**\***p<0.02, **p<0.002, ***p<0.0002).

Analysis of intestines collected from EW and RRV infected mice at 1 dpi by immunoblotting showed that compared to either mock or EW infection, RRV induces significant phosphorylation of STAT1 at Y701 (Fig. 5C), which is a convergent signaling response to different types of IFNs in the intestinal epithelium^15^. Collectively, these findings are consistent with the conclusion that, in contrast to homologous murine RV, heterologous simian RV strongly activates IFN-dependent STAT1 and pro-inflammatory responses in the small bowel.

### STAT1 activation status regulates the cleavage of multiple intestinal caspases during homologous vs. heterologous RV infection *in vivo*

Different IFNs, as well as several IFN-stimulated cytokines (such as IP10 and RANTES), induce caspase-regulated cell death to potentiate inflammation^26,27^. Differences between EW and RRV strains in their ability to regulate pSTAT1 and cytokine expression prompted us to examine the status of intestinal caspases following infection of suckling mice with these two RV strains. Using immunoblotting analysis, the cleavage status of the initiator caspases 8 and 9, and the executioner caspase 3, were examined in the small intestines of WT and STAT1^−/−^ mice infected with either EW or RRV RV (Fig. 6A). WT mice infected with RRV RV for 24 h undergo substantially higher levels of cleavage of caspases 8, 9, and 3 when compared to their mock-infected or EW RV counterparts (Fig. 6A-B), indicating that activation of both extrinsic and intrinsic intestinal apoptotic pathways is induced by heterologous RRV more efficiently than the homologous murine EW strain.

**Figure 6.**
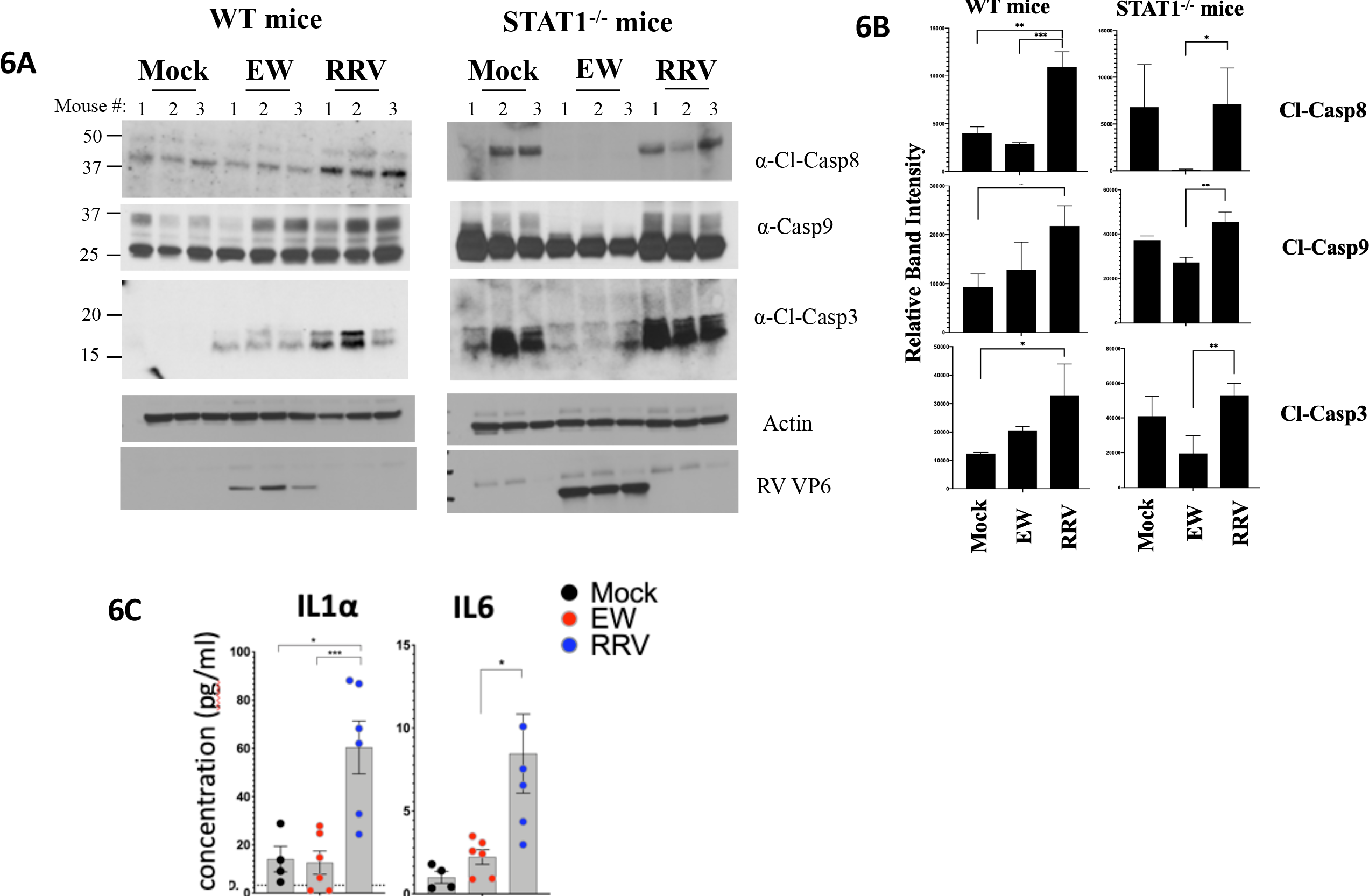
Homologous rotavirus inhibits the STAT1-mediated cleavage of multiple intestinal caspases. (A-B) Either wild type (WT) or STAT1-deficient mice were infected with EW or RRV RV strains (or mock infected). Twenty-four hours later, mice were sacrificed, and small intestinal lysates were analyzed by immunoblotting (A). Densitometric estimation of bands is shown in (B). (C) Mice infected as in (A) were sacrificed and intestines analyzed for the expression of cytokines by luminex-based assays. (**\***p<0.02, **p<0.002, ***p<0.0002)

Of note, the enhanced cleavage of caspases during RRV infection was dependent on the intestinal IFN response. Specifically, we found that differences in caspase 8, 9, and 3 cleavage between mock and RRV infected WT pups were abrogated following infection in STAT1 knockout mice (Fig. 6A-B). The cleavage of caspases during EW RV infection was also STAT1-mediated. However, EW RV-infected STAT1^−/−^ mice demonstrated cleaved caspases 8, 9, and 3 at levels much lower than in mock-infected controls or RRV infected mice (Fig. 6A and 6B), indicating that the basal cleavage efficiency of multiple caspases (i.e. occurring in the absence of infection) was impaired by homologous EW but not heterologous RRV RV.

Impaired caspase 8 cleavage can spontaneously induce acute intestinal inflammation through necroptosis, resulting in the secretion of the pro-inflammatory factor IL1α^28^. Therefore, we examined IL1α expression in the small intestine during EW and RRV infection (Fig. 6C). While the intestinal level of IL1α was no different between mock- and EW RV-infected suckling mice at 1 dpi, it was significantly elevated after infection with RRV RV^29^ (Fig. 6C). A similar difference was also observed for IL6, another common marker of intestinal inflammation^30^ (Fig. 6C). Thus, although murine EW RV inhibits caspase 8 cleavage when compared to simian RRV, it avoids inducing the pro-inflammatory cytokines IL1α or IL6 by a yet unknown mechanism. Collectively, these data identify an unrecognized role for STAT1 in mediating the cleavage of multiple intestinal caspases during RV infection. The findings also indicate that the ability of homologous RV to regulate IFN and STAT1 signaling *in vivo* (Figs. 4 and 5) likely dampens IFN regulated cell death and inflammation.

## DISCUSSION

Although they have been isolated from many different mammalian species, RVs are generally highly infectious only in their homologous host species. Robust RV replication occurs predominantly in the small intestinal mature villous tip epithelium and causes severe diarrhea, but only mild inflammation^6,7^. We and others demonstrated that both RV intestinal replication and accompanying inflammation are likely to be substantially regulated by host interferon (IFN) responses. Specifically, intestinal replication of a heterologous simian RRV strain can be significantly enhanced in suckling mice lacking IFNAR1, IFNGR1, and/or IFNLR1 receptors, or the convergent downstream transcription factor STAT1^3^. In the absence of STAT1, infection of suckling mice with the normally restricted RRV strain causes severe inflammation (mostly in the biliary tract), systemic spread, and mortality^8^. In contrast, replication of a homologous murine EW RV strain is quite resistant to IFNs^8^, and mice infected with EW RV are subsequently protected from lethal endotoxemia^15^, which is STAT1-mediated^24^. Paradoxically, EW RV robustly induces all three major IFN types^3,4^, raising the possibility that subsequent IFNR-directed antiviral and inflammatory signaling pathways are effectively blocked during homologous infection.

How homologous RV successfully inhibits the ability of intestinal IFNs to mediate antiviral and inflammatory effects is not yet well understood and was therefore examined in this study. We found that between 12-72 hpi, mice infected with murine EW RV exhibit sustained viral replication and increased transcription of the primary (i.e. induced prior to IFN amplification) murine type I IFN gene (IFNA4, Fig. 3A-B)^20^. However, transcriptional induction of the IFNL2/3, MX2, IFNG, RSAD2, IFIT2, and CXCL10 (Fig. 3A) genes in response to ongoing EW RV replication was considerably diminished by 72 hpi. Together, these data suggest that although ongoing EW RV replication triggers intestinal IFN induction signaling pathways, it progressively restricts intestinal IFN amplification responses. This conclusion is further supported by our finding that suckling mice infected with EW RV for 72 h are protected from severe small intestinal injury that normally occurs following systemic LPS exposure (Fig. 3B). This injury was reported to be driven primarily by STAT1 signaling in the mouse model of endotoxemia^24^. Using an alternate approach, we established that suckling mice infected with the STAT1-sensitive^3^ simian RRV strain exhibit significantly higher levels of intestinal ISG transcriptional induction (Fig. 5A), IFN-β protein expression (Fig. 5B), and STAT1-pY701 expression (Fig. 5C) compared to the homologous murine EW RV strain. In addition, expression of the IFN-stimulated pro-inflammatory cytokines IP10^31,32^ and RANTES^32–34^ are also higher in the intestines of heterologous simian RRV-infected mice (Fig. 5B). Collectively, these data support the conclusion that infection with homologous RV *in vivo* results in an inhibition of both IFN-mediated antiviral and inflammatory functions in the small intestine.

Caspase cleavage is a key parameter for determining both the magnitude and nature of inflammatory responses driven by IFNs and other cytokines^35,36^. Exposure to different IFNs can either induce caspase 8-mediated apoptosis^37^ or alternately, trigger inflammatory cell death (necroptosis) under circumstances when caspase 8 cleavage is impaired^38^. Data presented here (Fig. 6) demonstrate that murine EW RV infected mice express reduced cleavage of intestinal caspase 8, 9, and 3 when compared to RRV RV; cleavage levels being similar to the mock-infected control pups (Fig. 6A-B). In contrast, infection with RRV triggered the cleavage of all three caspases in the small intestine, indicating that both extrinsic and intrinsic apoptosis pathways are induced by the heterologous RRV despite its restricted replication capacity. Interestingly, activation of different caspases by RRV was contingent on the presence of STAT1, as differences in cleaved caspase 8, 9, and 3 expression between RRV and mock animals were eliminated in a STAT1 deficient background (Fig. 6A-B). The genetic KO of STAT1 in suckling mice infected with EW RV resulted in cleaved caspase levels lower than those in uninfected mock controls (Fig 6B). This was particularly evident for cleaved caspase 8 and 3 expression in the EW RV infected STAT1^−/−^ mice that were at or below the limit of detection (Fig. 6A and B). A likely explanation for these findings is that during the course of infection, EW RV regulation of IFN feedforward signaling (Fig. 4) and STAT1 phosphorylation (Fig. 5C) inhibits the efficiency of intestinal caspase 8 and 3 cleavage. Interestingly, inhibition of caspase 8 cleavage has been reported to induce spontaneous intestinal inflammation, primarily via necroptosis^28^. However, expression of IL-1α – a member of the IL-1 family that is released from necroptotic cells^39^, and of the pro-inflammatory cytokine IL-6, were not detected in EW RV (or mock infected control) pup intestines, but were significantly induced by RRV RV (Fig. 6C). Future studies will be required to understand how homologous RV inhibits the cleavage of intestinal caspases yet avoids causing intestinal inflammation. These findings nevertheless reveal a STAT1-dependent intestinal caspase activation pathway that is likely regulated by homologous RV to control inflammation.

Findings in this and earlier reports^3,4,8,9,15^ indicate that mechanisms by which RV regulates STAT1 signaling are likely critical determinants of rotaviral pathogenicity *in vivo*. We found earlier that several RVs (porcine, bovine, and simian RV strains) mediate the lysosomal degradation of multiple IFN receptor subunits (i.e. IFNAR1, IFNGR1, and IFNLR1) in RV VP6+ (i.e. infected) cells, whereas RV VP6− (i.e. bystander) cells expressed normal levels of the IFN receptors^15^. Remarkably, a porcine RV strain (SB1-A) was shown to also efficiently inhibit IFN-directed STAT1-Y701 phosphorylation in uninfected (i.e. VP6−) bystander cells^14^, indicating the existence of an alternative STAT1 inhibitory mechanism that operates beyond the virus-infected cell. In this study, we extended these findings and found that the simian RRV strain can also block IFN-β mediated STAT1-pY701 induction in both VP6-positive and -negative HT-29 cell populations (Fig. 1A-B). In addition, RRV infection prevented IFNβ-directed induction of the IRF7 protein in both infected and bystander cells (Fig. 1C). Together with our previous observation that both simian and porcine RV strains can mediate IFNR degradation^15^, these findings suggest that RV strategies to inhibit STAT1 signaling in infected and bystander cells are conserved across multiple strains that differ in the manner in which they regulate the IFN induction phase^17^.

Findings of this study also revealed a potential mechanism by which RV confers resistance to ectopic IFN stimulation to uninfected bystander cells^14^. Specifically, under infection conditions *in vitro* where only ~30% HT-29 cells were RRV VP6+, ectopic stimulation with IFN-I failed to evoke significant cellular transcriptional responses (Fig. 2). This inhibitory effect of RRV specifically targeted IFN signaling, as several of these measured transcripts could be efficiently induced during RRV infection by intracellular dsRNA stimulation (Fig. 2A). In addition, ectopic IFNAR1 stimulation during RRV infection also resulted in a significant decline of certain innate immune transcripts induced in response to RV (Fig. 2A). These collective findings not only demonstrate that RV infection can block IFNR-directed feedforward transcription but also suggest that RV represses transcription following ectopic IFN stimulation, which is characteristic of an IFNR-directed negative feedback transcription mode^2^.

The interesting possibility of negative feedback transcription effects is also supported from our studies in suckling mice which we previously showed that exogenous IFN I or II induced STAT1-Y701 in the small intestine with rapid kinetics (i.e. 30 min) when administered intra-peritoneally^15^. Remarkably, here we find that ectopic stimulation of either IFNR early during EW RV infection (i.e. at 12 hpi) substantially repressed several measured intestinal antiviral transcriptional responses (Fig. 4A). In particular, intestinal induction of both type I (*IFNB1* and *IFNA5*) and type III (*IL28*) IFN transcripts by EW RV was significantly dampened when IFNAR1 or IFNGR1 were ectopically stimulated. In addition, RV-induced MX2 transcription, which is reported to be primarily IFNLR1-driven in the intestinal epithelial compartment^23^, was eliminated following stimulation with either types I or II IFN.

Ectopic stimulation of IFNGR1 and IFNAR1 has been reported to elicit negative feedback signaling leading to IFN-resistance in the context of cancer and chronic viral infections^2,40^. The ability of homologous RV to elicit IFNAR1- and IFNGR1-stimulated feedback signaling suggests a potential mechanism underlying the ability of EW RV-infected mice to resist lethal endotoxemia^15^ and the accompanying intestinal damage (Fig. 3C). It will be interesting to identify the RV-mediated effectors involved in the broad suppression of multiple intestinal IFNR-dependent antiviral and inflammatory functions. As both type I and III IFNs were found to exert a profound antiviral activity against heterologous RV^3^, one possibility is that the endogenous induction of these IFNs during homologous EW RV infection^3,4,19^ serves to dampen intestinal antiviral, apoptotic, and inflammatory transcription and promote efficient viral replication.

## ACKNOWLEDGEMENTS

This work was supported by grants 1RO1 AI021362 (HBG), R56 AI125249 (HBG), and 1RO1 AI1125249 (HBG) from the National Institutes of Health, and 1IO 1BX000158-01A1 from the Veterans Affairs (HBG).

